# Effect of storage conditions on peripheral leukocytes transcriptome

**DOI:** 10.1101/2020.04.21.054296

**Authors:** Yanru Xing, Xi Yang, Haixiao Chen, Sujun Zhu, Jinjin Xu, Yuan Chen, Juan Zeng, Fang Chen, Mark Richard Johnson, Hui Jiang, Wen-Jing Wang

## Abstract

**Introduction:** Peripheral blood leukocytes are essential components of the innate and adaptive immune responses. Their transcriptome reflects an individual’s physiological and pathological state, consequently bulk and single-cell RNA sequencing have been used to assess health and disease. As RNA is dynamic and could be affected by *ex vivo* conditions before RNA stabilization, we have assessed the influence of temporary storage on the transcriptome.

**Methods:** We collected peripheral blood from six healthy donors and processed it immediately or stored it either at 4□ or room temperature (RT, 18–22□) for 2h, 6h and 24h. Total cellular RNA was extracted from leukocytes after red blood cells lysis and the transcriptome analyzed using RNA sequencing.

**Results:** We identified 152 up-regulated and 12 down-regulated coding genes in samples stored at 4□ for 24h with most of the up-regulated genes related to nucleosome assembly. More coding genes changed expression at RT, with 1218 increased and 1480 decreased. The increased genes were particularly related to mRNA processing and apoptosis, while the decreased genes were associated with neutrophil activation and cytokine production, implying a reduced proportion of neutrophils, which was confirmed by leukocyte subsets analysis. Most house-keeping genes changed expression during storage, but genes such as TUBB and C1orf43 were relatively stable and could serve as reference genes.

**Conclusion:** Temporary storage conditions profoundly affect leukocyte gene expression profiles and change leukocyte subset proportions. Blood samples stored at 4□ for 6h largely maintain their original transcriptome.

## Introduction

Leukocytes are important components of the peripheral immune system and play an essential role in protecting the body from infection. With the development of next-generation sequencing techniques, bulk RNA sequencing (RNA-seq) and single-cell RNA-seq have become increasingly popular ways to study the activity and function of leukocytes [1]. Bulk RNA-seq has revealed marked changes in the leukocyte transcriptome in different diseases [2-4] and single-cell RNA-seq has not only identified new subtypes of leukocytes [5, 6] but also found unique inflammatory gene expression profiles [7] in specific diseases. Thus, both bulk and single-cell leukocyte RNA-seq have established themselves as valuable tools in health research.

Although RNA is dynamic, it can be stabilized and stored at −80 in TRIzol Reagent for a long time before analysis, providing a convenient solution for large sample collections [8]. However, usually, leukocytes can not be isolated from blood immediately before TRIzol treatment, which leads to unpredictable changes in the transcriptome. Given the inherent challenges of blood collection tubes, leukocytes are susceptible to metabolic stress due to the lack of glucose and oxygen [9, 10]. As a result, the expression of hundreds of genes may be inadvertently changed. To obtain accurate leukocyte transcriptome, the first step should be to avoid or mitigate *ex vivo* changes on RNA [11]. PAXgene Blood RNA system can stabilize blood RNA immediately after taking blood [12], but single cells and plasma cannot be separated for further analysis. Blood collected in EDTA tube can be used as a liquid biopsy, including bulk and single-cell analysis, but little is known about the influence of temporary storage conditions on the leukocyte transcriptome. Understanding the impact of storage conditions on the transcriptome is critical for researchers to obtain meaningful and reproducible results. To optimize leukocyte transcriptome assessment we subjected peripheral blood to different short-term storage conditions, 4□ or room temperature (RT) for various periods, and assessed the influence on the leukocyte transcriptome.

## Materials and Methods

### Sample collection

Peripheral blood samples were obtained from three healthy women and three healthy men and were processed immediately or stored either at RT or 4□ for 2h, 6h and 24h (Fig. 1A). Leukocytes were subsequently isolated for RNA-seq.

**Fig. 1:**
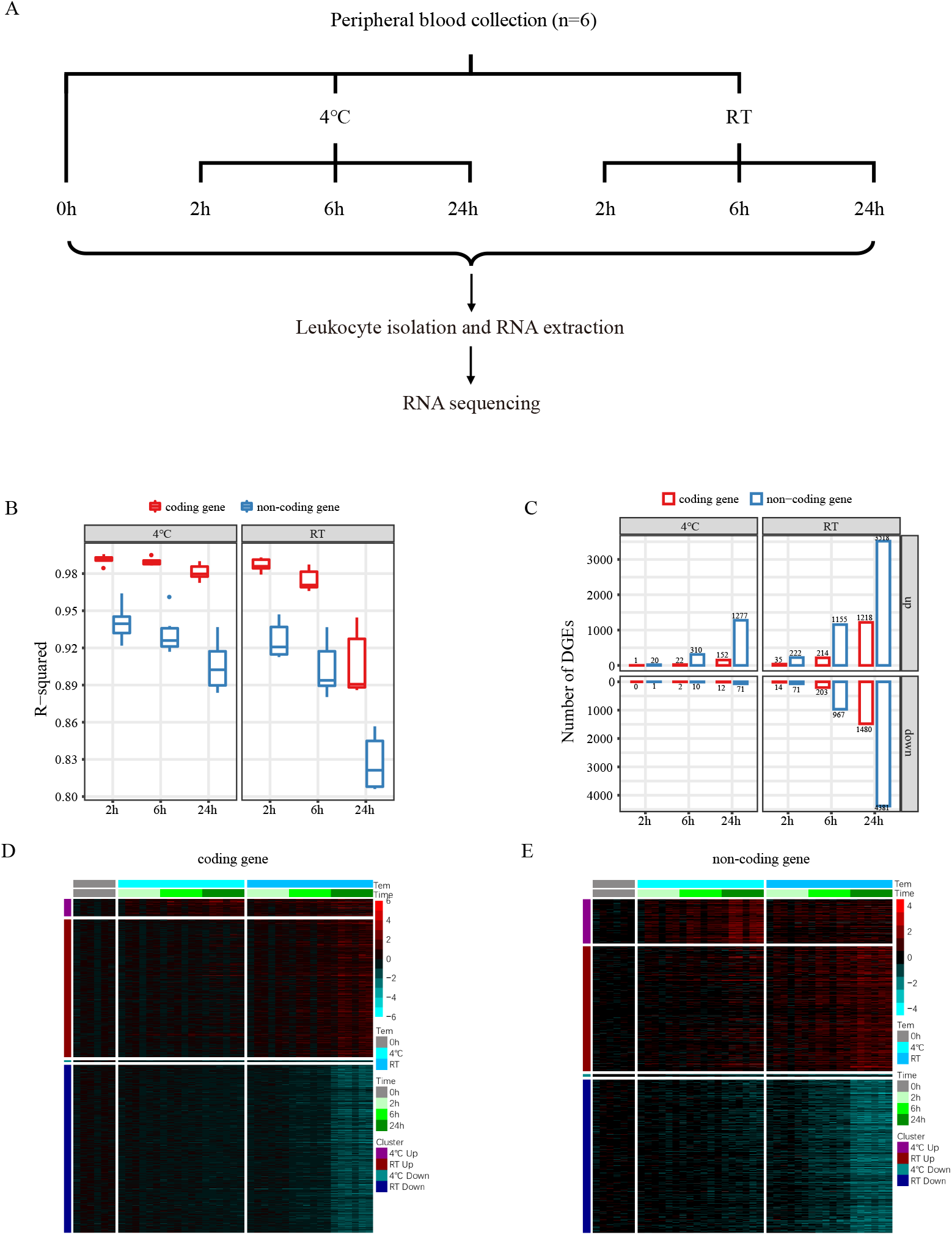
(A) Schematic of samples processing protocol: Whole blood was collected in EDTA tubes. Leukocyte isolated, RNA extracted and sequenced after stored in different conditions. (B) R-square between samples under different storage conditions and samples treated immediately. (C) Numbers of DEGs between samples under different storage conditions and samples treated immediately. (D, E) Heatmap represented different expressed coding and non-coding genes under different storage conditions. All gene expressions were transformed to log2(TPM+1)-mean0h log2(TPM+1).

### RNA isolation and sequencing

RNA from leukocytes was extracted by the TRIzol Reagent (Thermo Fisher, 10296028) following standard procedures as previously described [13]. RNA concentration and integrity were measured and calculated by Agilent 2100 bioanalyzer (Agilent Technologies, G2939A) and accompanying software. All RNA samples were with amounts greater than 200ng and RIN values of greater than 6 underwent library construction. Libraries were built using RNase H method [14] and sequenced on BGISEQ-500RS (single-end 50bp).

### Data preprocessing

Ribosomal ribonucleic acid (rRNA) was removed by SOAP2 (Version 2.21t) [15] and reads with low quality and adaptors was filtered by SOAPnuke (Version 1.5.6) [16] from raw data based on default parameters. Clean data were obtained in the FASTQ format. Kallisto [17] was used to align all clean reads to transcriptome references and obtain estimated reads counts and transcripts per million mapped reads (TPM) results. Transcriptome references were the refMrna.fa.gz file from the UCSC database with the removal of NR_RNA [18] and the NONCODEv5_human.fa.gz file from the NONCODE database [19].

### Differentially expressed gene (DEG) analysis

We used edgeR [20] and QL F-test to identify DEGs between samples stored under different conditions. Genes with count per million (CPM) greater than 1 in at least six samples were retained. Trimmed mean of M-values (TMM) [21] between each pair of samples was performed to minimize the log-fold changes between the samples for most genes. Q-values are calculated using the Benjamini-Hochberg [22] procedure to account for multiple testing. Genes performed Q-values ≤ 0.01 and fold change ≥ 2 between two groups were considered as DEGs.

### Weighted Gene Co-expression Network Analysis (WGCNA)

We merged all DEGs and retained all samples’ TPM data of these genes for WGCNA [23]. First, we used the scale-free topology criterion to choose the soft threshold β that R^2^>0.8 and the adjacency was transformed into topological overlap matrix (TOM) using a power function: a_mn_ = |r_mn_|^β^ (r_mn_: Pearson correlation coefficient for gene m and n). Second, the hierarchical clustering was used to cluster the topological overlap distance calculated by the adjacency matrix based on TOM dissimilarity measure with a minimum gene number is 50 and a_mn_ larger than 0.95. A module eigengene distance threshold of 0.25 was used to merge highly similar modules. Finally, we calculated the Pearson correlation between module eigengene [24] and storage time.

### Functional analysis

The biological process of coding genes in modules was performed by Metascape [25]. Top 5 clusters with their representative enriched terms (one per cluster) in each gene module were presented. Genes were annotated based on information in the GeneCards [26] and the Ensembl database [27]. The expression of house-keeping genes [28] under different storage conditions was described.

### Leukocyte subsets analysis and statistical analysis

The proportion of leukocyte subtypes was estimated by CIBERSORT [29] according to coding gene TPM. We used paired two-sided t-test to compare the differences between samples stored under different conditions and processed immediately, and *p* < 0.05 was considered to be significant. Data were presented as mean ± SD.

## Results

### Transcriptome influenced by storage conditions

RNA was successfully extracted from all samples, and the quality of RNA didn’t show significant differences. About 100M clean reads were obtained from each sample for further transcriptome analysis. The correlation of gene expression with samples processed immediately decreased with time and the expression profiles were more stable at 4□ than at RT (Fig. 1B). More coding and long non-coding genes changed expression at RT than at 4□, and the number of DEGs increased with time (Fig. 1C, 1D). Heatmap also showed that the expression of DEGs changed more at RT than 4□, this affected coding genes or non-coding genes similarly (Fig 1D, 1F).

### Storage time-associated co-expression modules

According to the expression of DEGs from 4□ and RT, the WGCNA successfully identified three (Fig. 2A) and four (Fig. 2B) gene integrative modules respectively. Further, we found the green and brown modules showed negative correlations with storage time, while the blue and black modules showed positive correlations (Fig. 2C, 2D). Gene numbers in the different modules were counted (Fig. 2E, 2F). Samples stored at 4□ had 14 down-regulated and 152 up-regulated coding genes, while samples stored at RT had 1499 genes down regulated and 1186 genes up regulated (Fig. 2G, 2H).

**Fig. 2:**
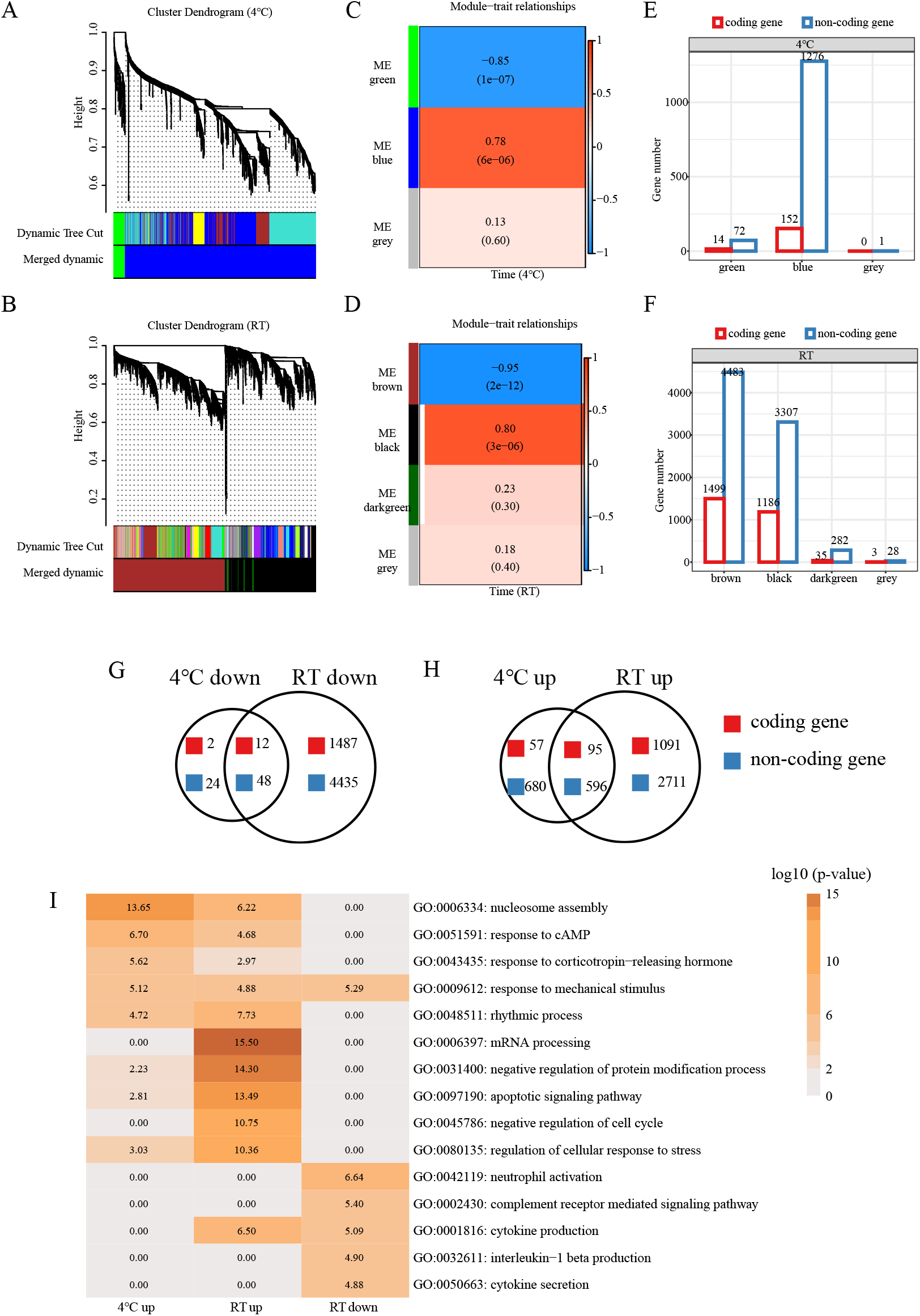
(A, B) Gene dendrogram by DEGs expression of 4□ and RT. The color row underneath the dendrogram showed the module assignment determined by the dynamic tree cut. (C, D) Heatmap of the correlation between module eigengenes and the storage time. Blue color represented a negative correlation, and red represented a positive correlation. (E, F) Coding and non-coding gene numbers in different modules. (G, H) Venn diagram showed the overlap of down/up-regulated genes between RT and 4□ within 24h. (I) Comparison of the biology process of up-regulated genes at 4□, RT and down-regulated genes at RT. Top 5 clusters with their representative enriched terms (one per cluster) in each gene module were presented (Methods).

### Biological process of coding genes in different modules

Coding genes for biological processes of nucleosome assembly and response to cAMP were up-regulated at 4□ and were also enriched to a lesser extent at RT (Fig. 2I). Coding genes up-regulated at RT were specifically associated with mRNA processing, negative regulation of protein modification process, apoptotic signaling pathway and negative regulation of cell cycle (Fig. 2H). While down-regulated coding genes at RT were particularly enriched for leukocytes activation processes in the immune response, such as neutrophil activation, complement receptor-mediated signaling pathway, cytokine production, interleukin-1 beta production and cell activation involved in immune response (Fig. 2H).

### The change of leukocyte subsets

The proportions of different leukocyte subsets varied in different storage conditions (Fig. 3A). Most subsets had constant relative proportions when samples stored at 4□ for 24h. However, the proportions of neutrophil, memory B cell and naïve CD4 T cell significantly decreased and CD8 T cell, resting memory CD4 T cell, regulatory T cell, and resting mast cell increased in samples stored at RT for 6h or 24h (Fig. 3B).

**Fig. 3:**
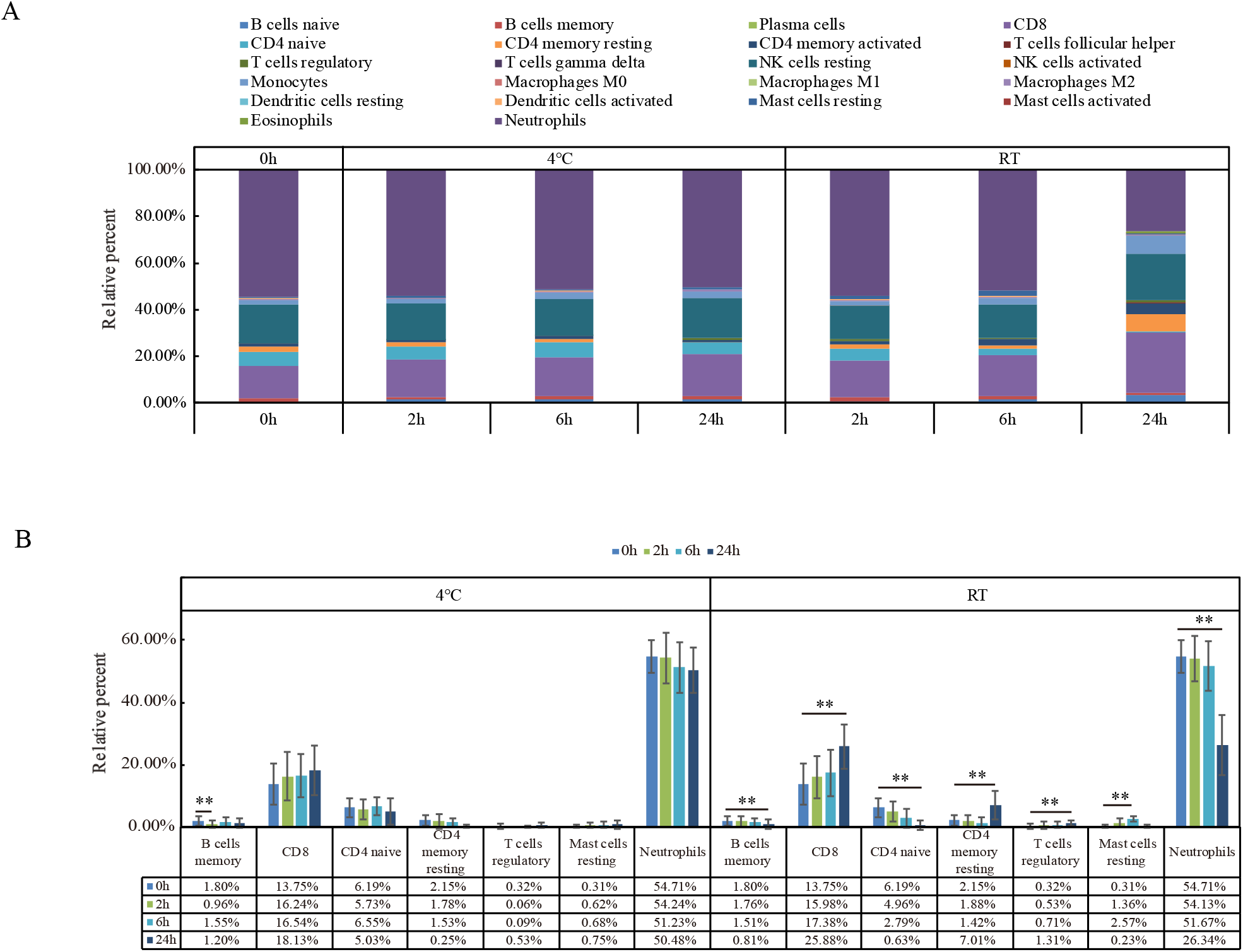
Leukocyte subsets analysis. (A) The relative percent of different cell subsets in different storage conditions. (B) Comparison of the relative percent of B cells memory, CD8, CD4 naive, CD4 memory resting, T cells regulatory, mast cells resting and neutrophils between samples under different storage conditions and samples treated immediately. * means p-value <0.05, ** means p-value <0.01.

### House-keeping genes expression during storage

The expression of C1orf43, TUBB, CHMP2A, SNRPD3, and RAB7A were relative stable both at RT and 4□ within 24h, while HSP90AA1, PSMB4, VCP, and REEP5 were significantly changed in samples stored for 24h both at 4□ or RT. The expression of the rest of house-keeping genes was relatively stable in samples stored at 4 (Fig. 4A). We identified 17 differentially expressed genes between females and males under different storage conditions. Most DEGs expression was more consistent at 4□ than at RT. The expression of such DEGs located in Y-chromosome and ZFP57 was independent of the storage conditions, while the expression of some genes was only significantly different between females and males when the samples stored at RT for 24h, such as IL1RN and IL-1B (Fig. 4B).

**Fig. 4:**
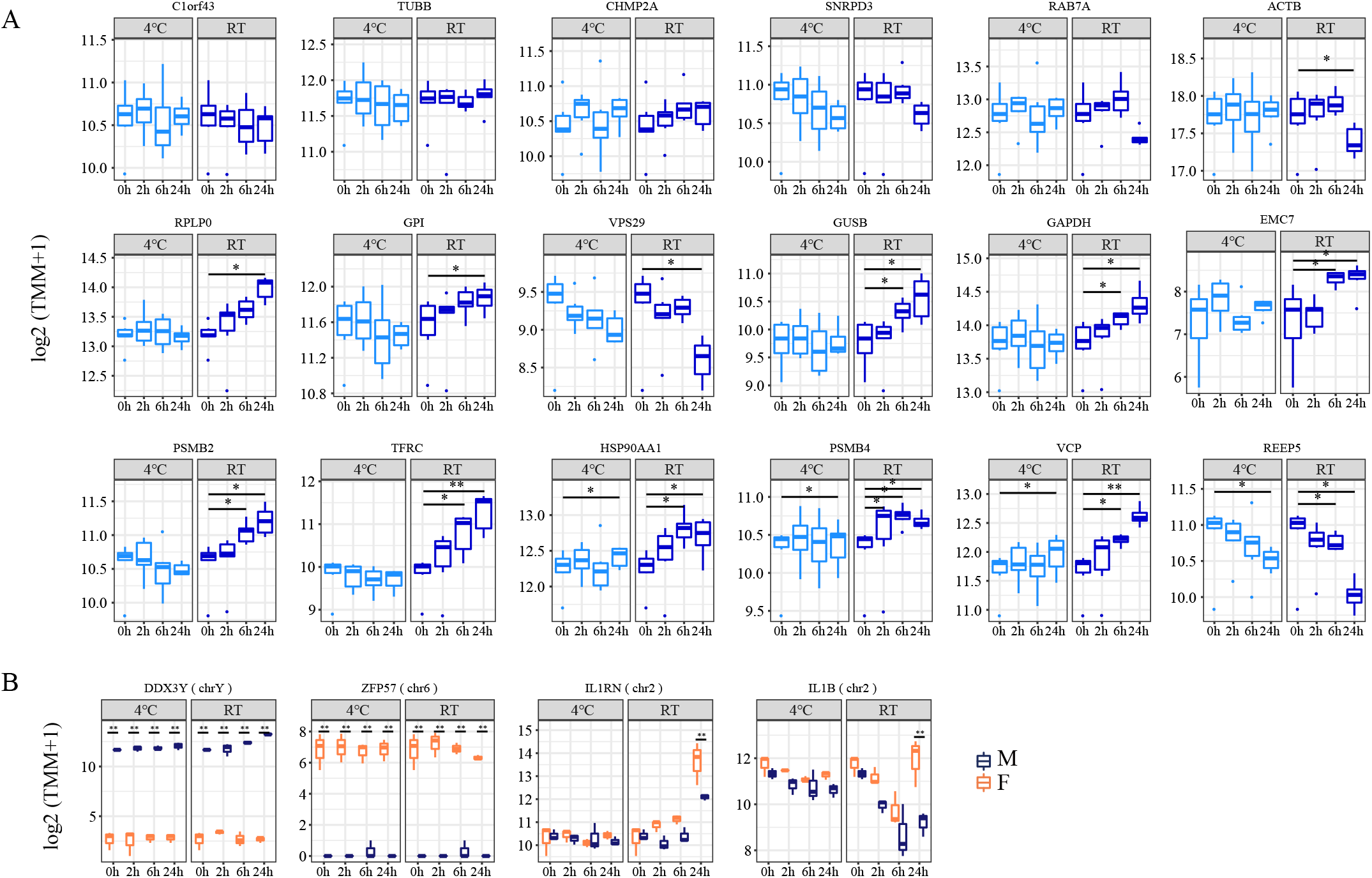
Expression of certain genes under different storage conditions. (A) Expression of house-keeping genes under different storage conditions. * means p-value <0.01, ** means p-value <0.01& fold change > 1. (B) Expression of DDX3Y, ZFP57, IL1RN and IL1B under different storage conditions. These genes performed different expression in genders. ** means p-adjust < 0.01 & fold change > 1. Gene expressions was transformed to log2(TMM+1).

## Discussion

How the mode of transport of peripheral blood samples between the clinic and the laboratory before RNA stabilization could affect bulk and single-cell RNA-seq of leukocytes remains unclear. Artificial changes, unrelated to the study, could materially affect the results [30, 31]. Thus, we investigated the effects of storage conditions on the leukocyte transcriptome by comparing peripheral blood samples stored at 4°C or RT with those processed immediately. The transcriptome changed more dramatically at RT than 4 *°C*, showing more up-regulated and down-regulated genes and variable leukocyte subsets proportions. Blood stored at 4°C and processed within 6h largely retains its original status, suggesting that samples should be stored at 4°C during transport.

Low temperatures can reduce metabolism, and previous studies have shown that RNA is relatively stable at 4°C [32]. From our data, it can be seen that the expression of most genes is relatively stable in samples stored at 4°C and that only 0.88% (166/18777) of coding genes and 1.66% (1349/81226) of non-coding genes were significantly influenced. Among the differentially expressed coding genes, 152 coding genes are up-regulated, and these genes were mostly enriched for nucleosome assembly. Previous studies reported that as the temperature increases, nucleosome DNA denatured [33] and our data indicate that low temperatures might repress transcription by altering the structure of chromatin. It is suggested that in studies designed to investigate the structure of nucleosomes and related functions, sample storage temperature should be taken into account.

Storing at RT within 24h did not influence RNA integrity of leukocytes, but significantly affected transcriptomic profiles, with the gene expression of 14.50% (2723/18777) of the coding genes and 9.97% (8100/81226) of the non-coding genes being significantly changed. According to the biology process of up and down-regulated coding genes, leukocytes stored at RT possibly underwent apoptosis and neutrophils activation. Blood cells preserved at EDTA tubes are under glucose and hypoxia deprivation that promotes cascades of biological alterations, causing inflammation [34, 35] and triggering neutrophils apoptosis [36-38]. As RT has a major impact on leukocyte transcriptome, the influence of temperature should not be ignored when dealing with samples stored at such a condition.

It was inferred that apoptosis happened when blood was stored at RT, so we used CIBERSORT to explore the leukocyte subsets proportions of each sample. We found that the relative proportion of neutrophils significantly decreased in samples stored at RT for 24h, while the leukocyte subsets composition was relatively stable at 4□ within 24h. As we know, neutrophils are the most abundant cell type in leukocytes [39, 40]. They have a short half-life (7−12 h *in vivo*) in peripheral blood, and are generally functionally quiescent under normal conditions [41]. The *ex vivo* temporary storage at 4 probably delays neutrophil apoptosis. Different cell types might have distinct fates under the same storage condition, leading to biased subsets composition. Therefore, the influence of storage conditions on identifying new cell types or evaluating cell type compositions especially in single-cell sequencing should be taken into consideration.

House-keeping genes are ideally expressed at relatively constant levels in most situations and critically important to our ability to identify changes in gene expression [28]. However, a previous study [42] showed that the expression of house-keeping genes could be altered by storage conditions, and that caution must be exercised when selecting genes for reference purposes. According to our results, C1orf43, TUBB, CHMP2A, SNRPD3, and RAB7A could be used as reference genes for leukocyte samples. IL1RN is associated with many autoimmune diseases like rheumatoid arthritis [43], which has a high incidence in women. Our study showed that IL1RN was not differentially expressed between males and females in the samples processed immediately, but its expression was greater in females than males after 24h storage at RT, which may indicate sex differences in the inflammatory response [44]. It is suggested that gene expression is not only affected by *ex vivo* conditions but also influenced by the genetic background, consequently long-term storage before RNA stabilization could markedly alter the transcriptomic profile.

## Conclusion

Peripheral blood collected into EDTA tubes, stored at 4□ and processed within 6h largely maintain the original transcriptome. Temporary storage of blood samples at different temperatures and for longer periods could change the gene expression profile and leukocyte subset proportions. C1orf43, TUBB, CHMP2A, SNRPD3, and RAB7A could serve as reference genes under such conditions. It is critically important to consider how sample processing and storage could influence bulk and single-cell RNA-seq to optimize the results of transcriptomic analysis.

## Acknowledgments

We are grateful to all healthy volunteers who donating blood for this study. We also thank Dr Qing Zhou and Dr Zhongzhen Liu in our team for sharing their advice on this study.

## Compliance with Ethical Standards

### Funding

This project is supported by the National Key Research and Development Program of China (No.2018YFC1004900), the National Natural Science Foundation of China (No.81300075), the Science, Technology and Innovation Commission of Shenzhen Municipality under grant (No. JCYJ20170412152854656, JCYJ20180703093402288).

### Informed Consent

All volunteers signed informed consent, and the study protocol was approved by the BGI Institutional Review Board (NO. BGI-IRB 17166).

### Availability of data

Raw RNA-seq data have been submitted to the CNGB Nucleotide Sequence Archive (CNSA: https://db.cngb.org/cnsa; accession number CNP0000266). Data generated or analyzed during this study are included in this published article.

